# Ongoing lymphoid HIV production drives pyroptosis and GLP-1 counter-regulation in ART-suppressed infection

**DOI:** 10.64898/2026.01.09.698696

**Authors:** Peter A. Crawford, Joshua Rhein, Jeffrey G. Chipman, Gregory J. Beilman, Ross Cromarty, Kevin Escandón, Jodi Anderson, Garritt Wieking, Andrew Johnston, Afeefa Ahmed, Jarrett Reichel, Alexander Khoruts, Christopher M. Basting, Nataliia Kuchma, Jason V. Baker, Nichole R. Klatt, Ashley T. Haase, Timothy W. Schacker

## Abstract

Despite effective antiretroviral therapy (ART), many people with HIV (PWH) exhibit persistent immune activation (IA) and suffer metabolic comorbidities. We investigated whether residual HIV production in lymphoid tissues drives IA. Among 20 ART-suppressed PWH, HIV RNA^+^ cells were detected in lymph nodes and correlated directly with markers of pyroptosis, assessed via cleaved gasdermin D positivity, but not with most plasma cytokines or IA markers. Notably, glucagon-like peptide 1 (GLP-1), an enteroendocrine hormone with anti-inflammatory roles, was upregulated in the ileum of PWH and correlated directly with systemic cytokines but inversely with lymph node pyroptosis. These findings suggest that chronic occult inflammation in people with successfully suppressed HIV infection is mediated by persistent virus production in lymph nodes leading to pyroptosis, which may trigger compensatory anti-inflammatory enteroendocrine activation that may dampen pyroptosis. Targeting pyroptosis or enhancing GLP-1 signaling represent potential therapeutic strategies for modulating IA and managing metabolic comorbidities in PWH.

**Graphical abstract:** 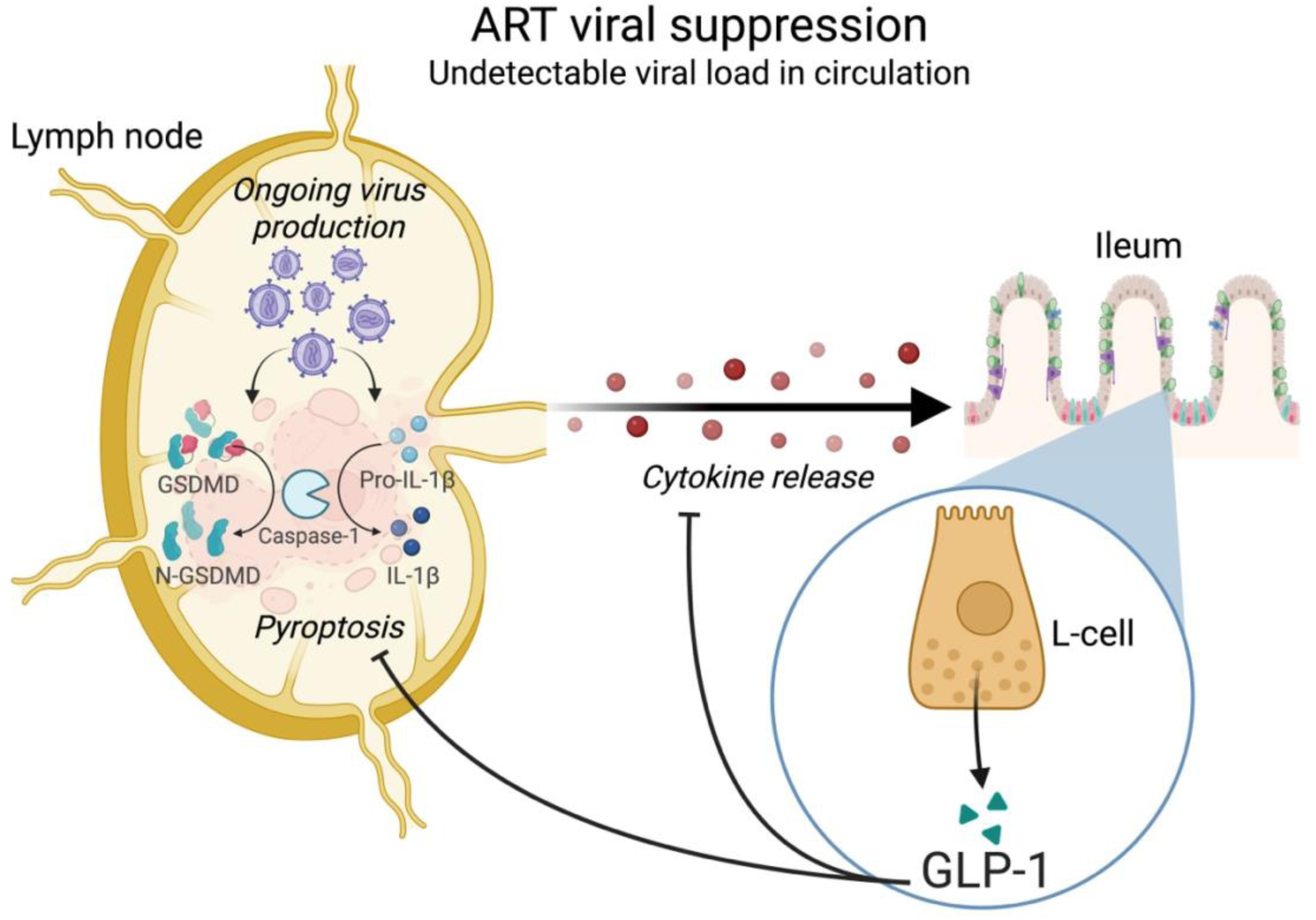

## Introduction

Antiretroviral therapy (ART) has markedly prolonged survival and mitigated morbidity for people living with HIV (PWH), however major challenges remain. Many individuals have evidence of persistent immune activation (IA), with levels of circulating proinflammatory cytokines significantly higher than those observed in HIV-negative people and increased numbers of activated T cells, especially CD8^+^ T cells^1,2^. Mechanisms underlying this state of persistent IA are unknown, but other chronic viral infections (e.g., those caused by herpes viruses) and microbial translocation are possible explanations^3–7^, as well as ongoing production of HIV in lymphatic tissues despite suppression of plasma viremia. We and others have consistently documented HIV-producing cells in lymph nodes (LNs) and gut despite long-term ART^8–12^. It is possible that this ongoing low-level HIV production in tissue sanctuaries triggers an immune response that maintains heightened levels of IA.

Finding the root cause of IA in PWH is important because, while prevalence can vary considerably across studies and populations, at least 35% of PWH controlled on ART develop metabolic complications that might be associated with sustained IA^13,14^. These include diabetes, fat redistribution, cardiovascular disease, and metabolic dysfunction-associated steatotic liver disease (MASLD)^15–17^. In the past, ART, especially nucleoside reverse transcriptase inhibitors and protease inhibitors, were linked to metabolic dysfunction, however new generation antiretroviral drugs do not carry the same risk for those metabolic abnormalities^18,19^. The cause of metabolic derangements in the modern ART era remains unknown but in HIV-negative individuals these metabolic conditions are often associated with chronic inflammation related to obesity^20^.

In this study we explore the relationships between (i) persistent virus production and IA and (ii) the relationship between IA and the expression of the incretin hormone glucagon-like peptide 1 (GLP-1). Our hypothesis was that persistent HIV RNA production in lymphatic tissues—though undetectable in plasma—leads to chronic inflammation and IA, resulting in abnormalities in the production of the incretin GLP-1 and other protective hormones, which, in turn may predispose to metabolic abnormalities increasingly seen in PWH, including diabetes, steatosis, and obesity.

## Results

### Cohort description

We recruited 20 people with chronic HIV infection on stable ART and with undetectable plasma viremia. The median age was 43.5 years, the median CD4^+^ T cell count was 667 cells/µl, and the median CD4/CD8 ratio was 0.82. Plasma HIV RNA levels were below the limit of detection for all study participants at the time of biopsy. Individuals had an inguinal LN biopsy and colonoscopy to collect gut-associated lymphoid tissue (GALT). In addition, we collected peripheral blood to measure proinflammatory cytokines and T cells in plasma. A total of 21 age-matched HIV-negative participants were recruited to obtain plasma for comparative analyses of proinflammatory cytokines. We also used existing LNs (*n* = 12) and ileum (*n* = 7) samples from HIV-negative participants enrolled in prior studies as negative controls. There were no body mass index (BMI) differences between PWH and HIV-negative controls. The demographic characteristics of all groups (PWH, HIV-negative individuals with plasma, and HIV-negative individuals with tissue samples) are shown in **Table 1**.

**Table 1.** Demographic and clinical characteristics of participants.

| Characteristic | PWH<br>(*n* = 20) | HIV-negative<br>participants<br>(plasma, *n* = 21) | HIV-negative<br>participants<br>(LN tissue, *n* = 12) | HIV-negative<br>participants<br>(ileum tissue, *n* = 7) |
| --- | --- | --- | --- | --- |
| Age, median<br>(IQR), years | 43.5 (32.0, 56.8) | 45.0 (34.0, 51.0) | 44.0 (28.3, 51.8) | 48.0 (33.0, 51.5) |
| Male sex at birth,<br>n (%) | 20 (100%) | 20 (95%) | 10 (83.3%) | 6 (85.7%) |
| Race, n (%) |  |  |  |  |
| White | 7 (35%) | 17 (81%) | 7 (58.3%) | 4 (57.1%) |
| Black | 7 (35%) | 0 (0%) | 1 (8.3%) | 2 (28.6%) |
| Other | 6 (30%) | 4 (19%) | 4 (33.3%) | 1 (14.3%) |
| Hispanic/Latino<br>ethnicity, n (%) | 3 (%) | 1 (4.8%) | 2 (16.7%) | 0 (0%) |
| BMI, median<br>(IQR), kg/m<sup>2</sup> | 25.9 (24.2, 28.1) | NA | 26.1 (20.4, 26.8)<sup>B</sup> | 27.1 (26.3, 28.3)<sup>C</sup> |
| Nadir CD4<sup>+</sup> T cell<br>count, median<br>(IQR), cells/μl | 296.0 (93.3, 407.8) | - | - | - |
| Time since HIV<br>diagnosis, median<br>(IQR), years | 9.5 (3.3, 17.0) | - | - | - |
| CD4<sup>+</sup> T cell<br>count<sup>A</sup>, median<br>(IQR), cells/μl | 667 (394, 810) | NA | 867 (715, 938)<sup>D</sup> | 921 (722, 954)<sup>E</sup> |
| CD8<sup>+</sup> T cell<br>count<sup>A</sup>, median<br>(IQR), cells/μl | 919.5 (613, 1222) | NA | 445 (313, 530)<sup>D</sup> | 375 (291, 500)<sup>E</sup> |
| CD4/CD8 ratio<sup>A</sup>,<br>median (IQR) | 0.82 (0.39, 1.14) | NA | 1.76 (1.34, 2.97)<sup>D</sup> | 1.84 (1.60, 3.65)<sup>E</sup> |

NA, not available. <sup>a</sup>Measured at the time of study procedures. <sup>b</sup>Available data ( $n = 10$ ). <sup>c</sup>Available data ( $n = 6$ ). <sup>d</sup>Available data ( $n = 11$ ). <sup>e</sup>Available data ( $n = 5$ ).

### Measures of persistent HIV RNA production in LNs

We used HIV RNA *in situ* hybridization (RNAscope) to measure the frequency of HIV-producing (vRNA^+^) cells in LNs. Collected samples were sufficient to measure the frequency of vRNA^+^ cells in 19 out of 20 PWH. HIV RNA^+^ cells were detected in all 19 LNs. The median log frequency (IQR) was 4.26 cells/g LN (3.81–4.49), which is the equivalent of approximately 30 vRNA^+^ cells/10^6^ CD4^+^ T cells. This is consistent with our prior reports of persistent virus production and overall frequency of HIV-infected cells in PWH while on ART^8,9,12,21,22^.

### Plasma cytokine measurements

We next measured levels of 13 plasma cytokines in the study PWH and the age-matched HIV-negative participants (**Fig. 1A–1H**). We found significant elevation of 7 of the plasma cytokines, specifically IL-1β, IL-2, IL-8, IL-12p70, IL-23, MIP-1β, and TNF-α, with no changes in IL-18, IFN-γ, IL-10, IL-17A, IL-5, or IL-6. This pattern of cytokine response is consistent with innate IA with antigen-presenting cell (APC)-derived proinflammatory cytokine induction (a core mechanism in metabolic syndrome pathogenesis), but no full adaptive T cell polarization or strong regulatory counterbalance.

**Fig. 1.**
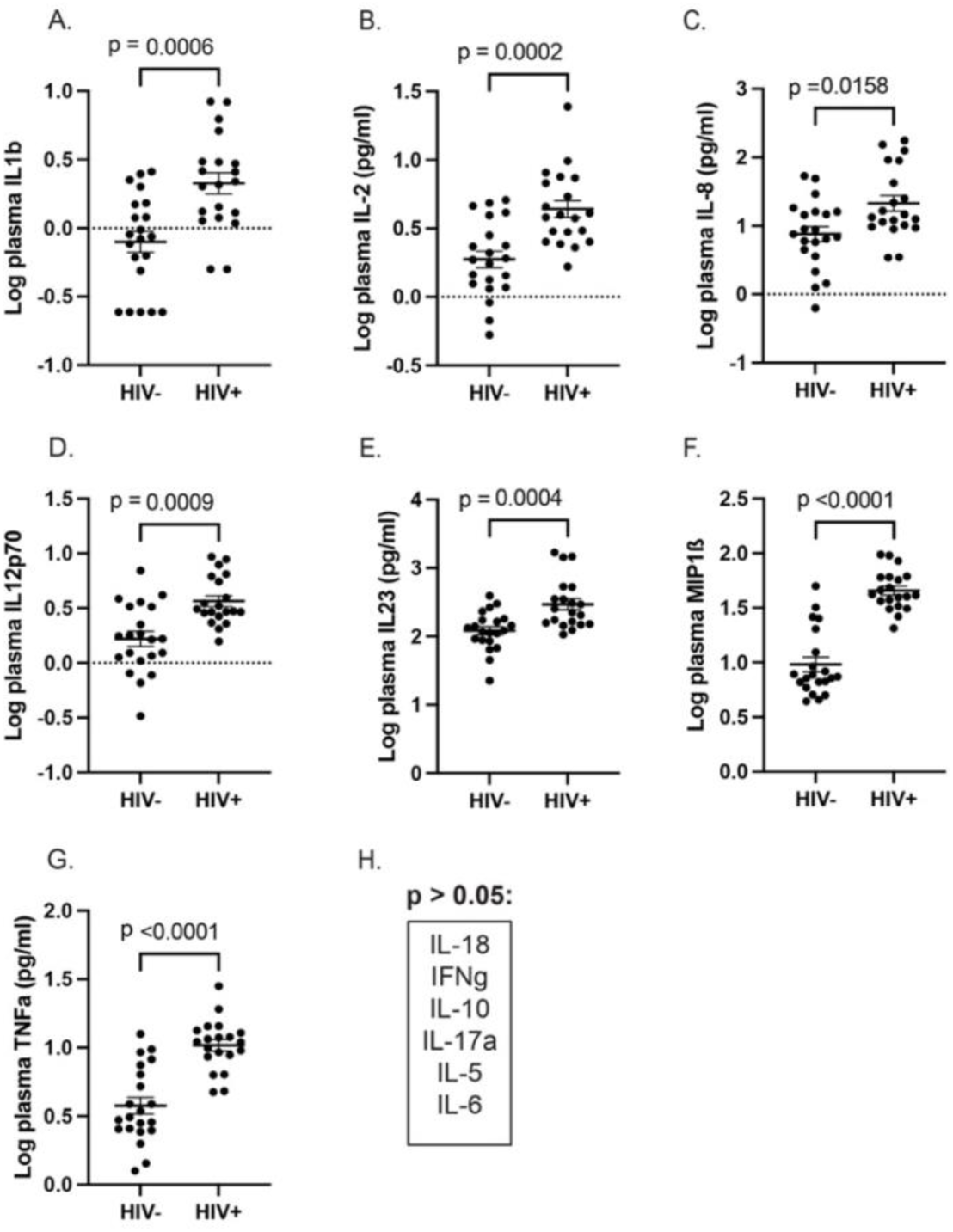
Increase in plasma cytokines in treated HIV infection. There was a significant elevation in the levels of plasma cytokines in the PWH compared to the HIV-negative group for: (**A**), IL-1β; (**B**), IL-2; (**C**), IL-8; (**D**), IL-12p70; (**E**), IL-23; (**F**), MIP-1β; and (**G**), TNF-α. (**H**), There was no difference in the plasma levels of IL-18, IFN-γ, IL-10, IL-17A, IL-5, or IL-6 between the groups.

### Relationship between persistent HIV RNA production in LNs and immune activation

We first compared the frequency of vRNA^+^ cells in LNs to plasma cytokine concentrations and found no correlation between vRNA^+^ cells in LNs and any plasma cytokine measured except IL-8 (**Fig. 2A**, *p* = 0.015, *r*^2^ = 0.315). Thus, ongoing HIV RNA production in LNs does not appear to be associated with these circulating biomarkers of immune activation. Analyses with *r*^2^ and *p* values for the remaining plasma cytokines are presented in **Supplementary Table 1**.

**Fig. 2.**
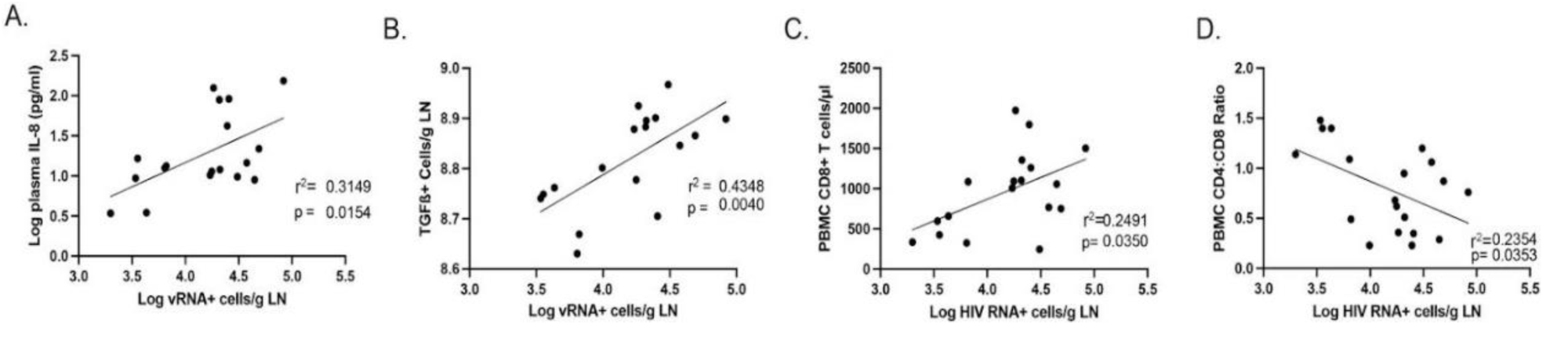
(**A**), Relationship between the frequency of vRNA^+^ cells in LNs and plasma IL-8. (**B**), Relationship between the frequency of vRNA^+^ cells and TGF-β^+^ cells in LNs. (**C**), Relationship between the frequency of vRNA^+^ cells and CD8^+^ T cells in peripheral blood. (**D**), Relationship between the frequency of vRNA^+^ cells and CD4/CD8 ratio in peripheral blood.

Next, we combined immunohistochemistry (IHC) with quantitative image analysis to determine the frequency of cells in the LN expressing IL-6 or TGF-β and the cell-associated IA markers CD25 and Ki-67. The median (IQR) frequency for cells expressing IL-6 was 7.86 cells/g LN (7.75–7.98) and for cells expressing TGF-β was 8.82 cells/g LN (8.71–8.90). The median (IQR) frequency for CD25^+^ cells was 7.92 cells/g LN (7.76–8.18), and for Ki-67, it was 8.32 cells/g LN (8.71–8.90). Frequencies of LN vRNA^+^ cells and TGF-β^+^ cells were directly correlated (**Fig. 2B**, *p* = 0.004, *r*^2^ = 0.438) but LN vRNA^+^ cells were not correlated with the frequency of IL-6^+^, CD25^+^, or Ki-67^+^ cells. These data suggest that ongoing virus production in LNs does not appear to be associated with LN cells expressing common IA markers or IL-6 but is associated with expression of TGF-β.

Finally, we examined the relationship between the frequency of vRNA^+^ cells in LNs and biomarkers of immune reconstitution in PBMCs. The size of the CD8^+^ T cell population and the CD4/CD8 ratio have been used as surrogate markers of immunologic recovery in PWH following ART initiation. We found that the higher the frequency of vRNA^+^ cells in LNs, the larger the population of CD8^+^ T cells in peripheral blood (**Fig. 2C**, *p* = 0.035, *r*^2^ = 0.249). In addition, the higher the frequency of vRNA^+^ cells in LNs, the lower the CD4/CD8 ratio (**Fig. 2D**, *p* = 0.0187, *r*^2^= 0.235).

### Pyroptosis in LNs

The lack of correlation between the frequency of HIV RNA-producing cells and measures of circulating proinflammatory cytokines or IA markers was unexpected because plasma levels of several proinflammatory cytokines were significantly elevated in PWH compared with HIV-negative individuals (**Fig. 1A-1G**). These results suggest that either ongoing HIV production is not driving IA in PWH or that a previously undetermined process originating within the LN drives chronic IA and inflammation. Plasma samples of PWH showed higher circulating IL-1β, which is increased in the presence of inflammasome activation and pyroptosis, an inflammatory form of programmed cell death associated with cell membrane rupture and release of proinflammatory cytokines^23,24^. This process has recently been shown to play a significant role in the pathogenesis of COVID-19 pneumonia^25^ and other viral infections. Therefore, we reasoned that a pyroptotic cascade may contribute to immune activation pathogenesis in treated HIV infection. To test this hypothesis, LN sections were incubated with antibodies directed against cleaved gasdermin D (GSD), a pyroptotic protein that when cleaved by caspase 1 or caspase-4/5, forms pores in the cell membrane, resulting in the release of proinflammatory cytokines from macrophages and other cells of the innate immune system recruited to the area (**Fig. 3A**)^23^. In addition, we stained tissues with antibodies against IL-18, demonstrating that all GSD^+^ cells were also IL-18^+^, a cytokine implicated in the pathogenesis of pyroptosis. While macrophage clearance of apoptotic cells through efferocytosis is typically not associated with an inflammatory response, clearance of pyroptotic cells programs macrophages to release proinflammatory cytokines^26^. Indeed, a triple-label stain for cleaved GSD, IL-6, and the macrophage marker CD68 showed that GSD^+^ cells were consistently CD68^+^ and most were also IL-6^+^ (**Fig. 3B**). This high-resolution image (60x) demonstrates intact cells membranes of the CD68^+^ cells, which suggests phagocytosis of pyroptotic cells and release of proinflammatory cytokines. The plasma cytokine profile of the PWH (**Fig. 1A-1G**) supports this hypothesis as well. The median (IQR) log frequency of GSD^+^ cells was 6.46 cells/g LN (5.92–7.10), significantly higher than the median (IQR) log frequency of GSD^+^ cells in the 12 LNs from HIV-negative individuals, 5.77 cells/g LN (5.25–5.91) (**Fig. 3C**, *p* = 0.001, Mann-Whitney *U* test). Finally, the frequency of GSD^+^ cells was directly correlated with the frequency of vRNA^+^ cells (**Fig. 3D**, *r*^2^ = 0.34, *p* = 0.0218), suggesting that ongoing HIV RNA production in LNs is correlated with pyroptosis.

**Fig. 3.**
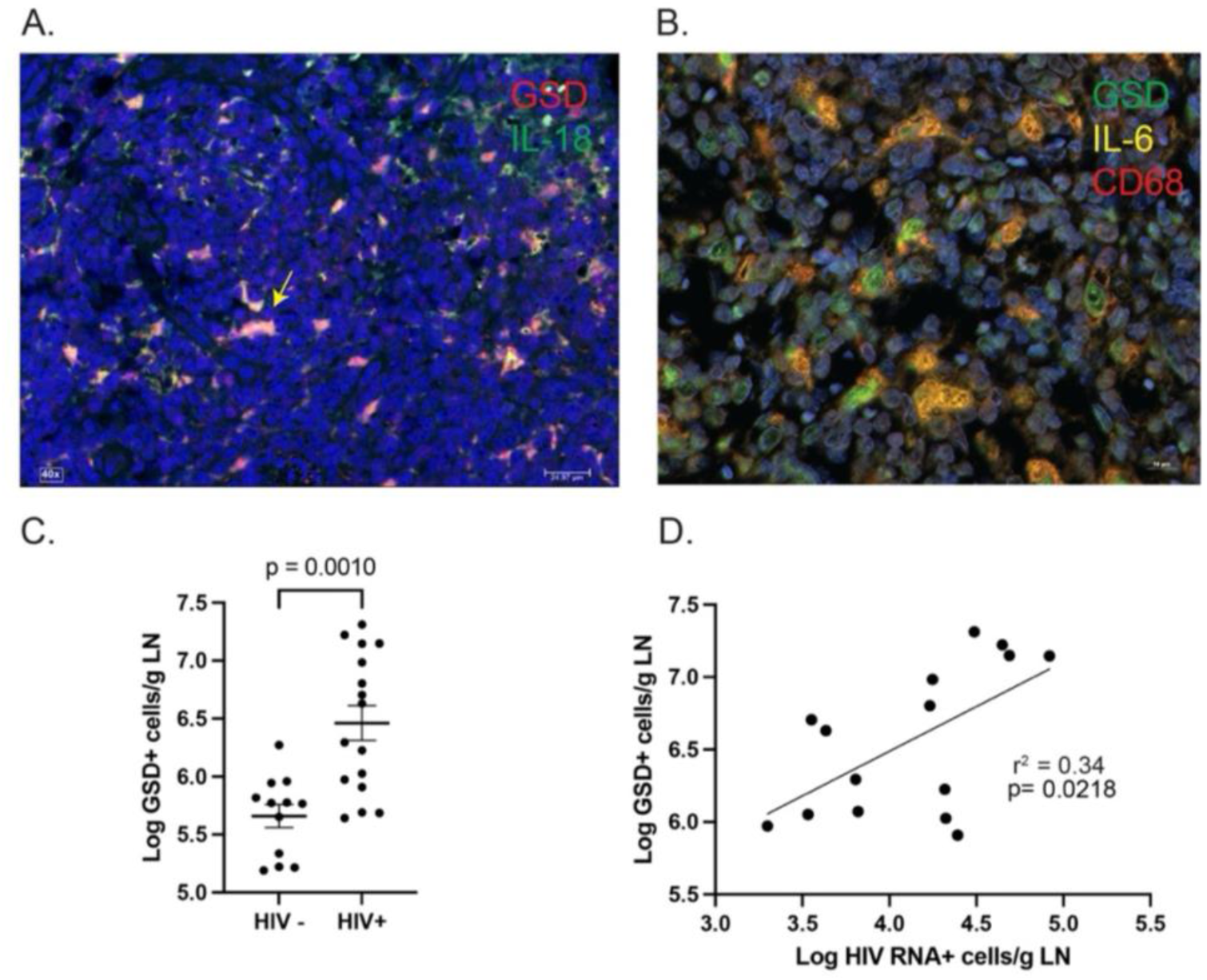
Pyroptosis in LNs. (**A**), Antibodies against cleaved GSD and IL-18 demonstrate the presence of pyroptotic cells that are GSD^+^ and IL-18^+^. GSD^+^/IL-18^+^ cells appear pink and the yellow arrow points to a representative double-positive cell. (**B**), We used antibodies directed against GSD, IL-6, and CD68 and found that most pyroptotic cells were CD68^+^ (i.e., macrophages) and were also IL-6^+^. This high-resolution image (60x) demonstrates that virtually all CD68^+^ macrophages that were GSD^+^ appeared intact, suggesting efferocytosis. (**C**), Quantitative image analyses demonstrated that GSD+ cells occur in LNs of PWH at ~1 log greater median frequency than HIV-negative individuals (*p* = 0.0010, Mann-Whitney *U* test). (**D**), The frequency of GSD^+^ cells in LNs of PWH correlated with the frequency of HIV RNA^+^ cells (*r*^2^ = 0.34, *p* = 0.0218).

### Pyroptosis and GLP-1

Pyroptosis is implicated in the pathogenesis of numerous diseases increasingly observed as co-morbidities in PWH, including lipodystrophy, fat depot redistribution, diabetes, MASLD, Alzheimer disease and other dementias^27–30^. Emerging multi-omics evidence suggests that inflammatory cell death could underlie cardiometabolic comorbidities, including PWH^20,31,32^. Therefore, we examined the relationship between the frequency GSD^+^ cells in the LN and the frequency of ileal cells expressing the gut enteroendocrine cell-derived incretin hormone GLP-1, an anti-inflammatory hormone that also improves glycemia and promotes feeding satiety^33^.

We measured the frequency of GLP-1^+^ cells in ileal samples of 28 PWH (the 20 participants described above plus 8 HIV^+^ participants from previous studies with similar demographics) and in 7 HIV-negative controls (**Fig. 4A**). The median (IQR) frequency of GLP-1^+^ cells/g ileum in HIV-negative individuals was 2.99 x 10^6^ cells/g ileum (1.76 x 10^6^–3.25 x 10^6^) and in the PWH it was 4.32 x 10^6^ cells/g ileum (2.53 x 10^6^–6.42 x 10^6^) (**Fig. 4B**) representing a statistically significant 36.4% increase in the PWH (*p* = 0.0095, Welch *t* test).

**Fig. 4.**
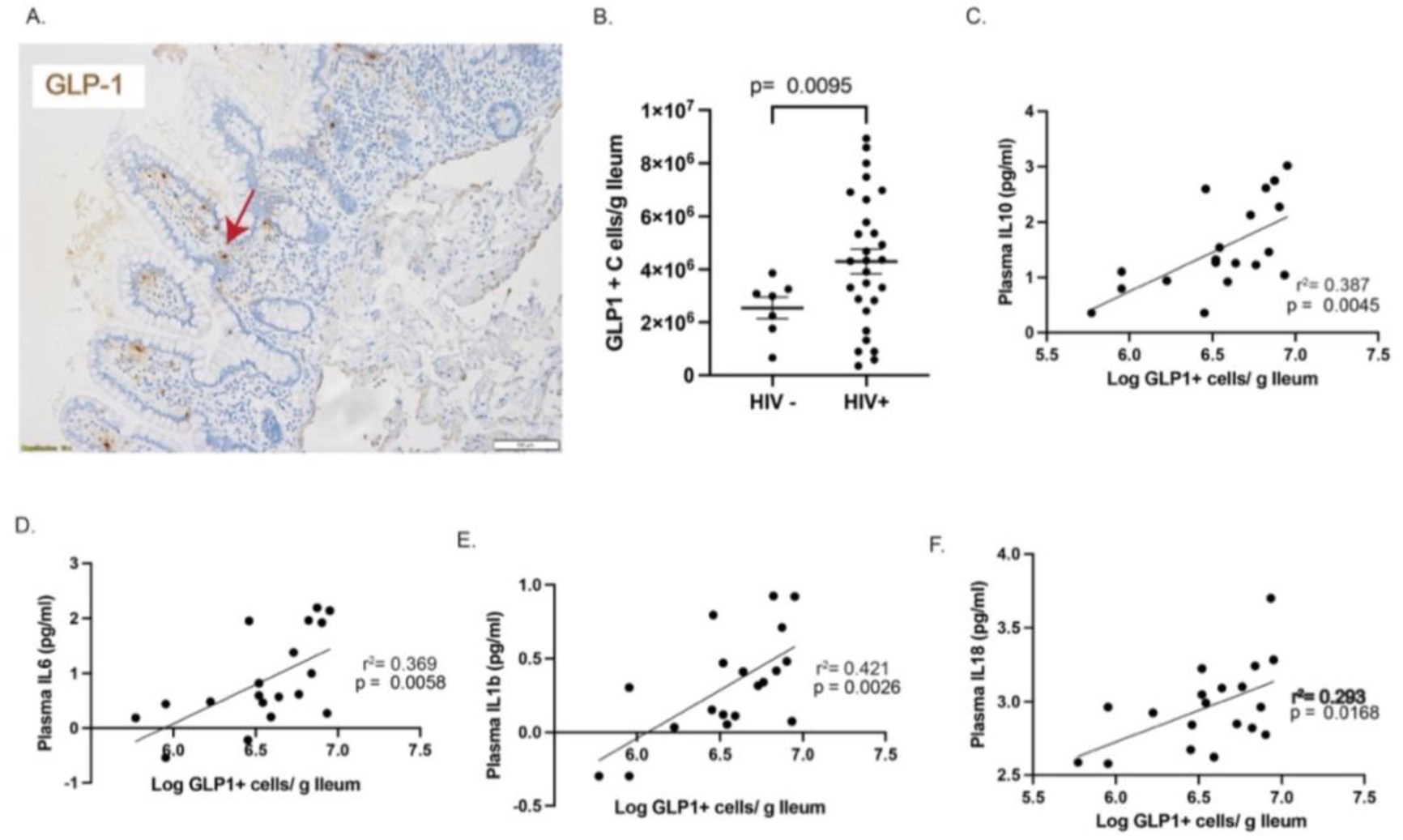
(**A**), Ileal tissues stained with antibodies against GLP-1. (**B**), Statistically significant increase in the frequency of GLP-1^+^ cells/g ileal tissue in PWH compared with HIV-negative participants. (**C**), The frequency of GLP-1^+^ cells directly correlated with plasma measures of IL-10. (**D**), The frequency of GLP-1^+^ cells directly correlated with plasma measures of IL-6. (**E**), The frequency of GLP-1^+^ cells directly correlated with plasma measures of IL-1β. **f,** The frequency of GLP-1^+^ cells directly correlated with plasma measures of IL-18.

Because incretins and incretin-mimetic therapeutic agents possess anti-inflammatory properties^33–37^, we next sought to determine the relationship between the frequency of GLP-1^+^ ileal cells and the circulating concentrations of cytokines that were augmented in PWH. We found significant relationships between frequency of GLP-1^+^ ileal cells and plasma measures of IL-10 (*r*^2^ = 0.387, *p* = 0.0045), IL-6 (*r*^2^ = 0.369, *p* = 0.0058), IL-1β (*r*^2^ = 0.421, *p* = 0.0026), and IL-18 (*r*^2^ = 0.293, *p* = 0.0168) (**Fig. 4C-4F**). Statistically significant direct correlations were observed between the frequency of GLP-1^+^ ileal cells and circulating concentrations of pyroptotic cytokines IL-10, IL-1β, IL-6, and IL-18 but not other cytokines (**Supplementary Table 2).**

Finally, we compared the frequency of GLP-1^+^ cells in the ileum to the frequencies of GSD^+^ cells and vRNA^+^ cells in the LNs. While no correlation between the frequency of vRNA^+^ cells in LNs and GLP-1^+^ cells in ileum was observed (**Fig. 5A**), the frequency of GSD^+^ cells in the LN and GLP-1^+^ cells in the ileum were inversely correlated (**Fig. 5B**, *p* = 0.0026, *r*^2^ = 0.51). Taken together, these results suggest that IA in PWH induces the production of the incretin hormone GLP-1 in the ileum, which in turn could serve to dampen pyroptosis and mitigate the pathogenesis of the infection during ART.

**Fig. 5.**
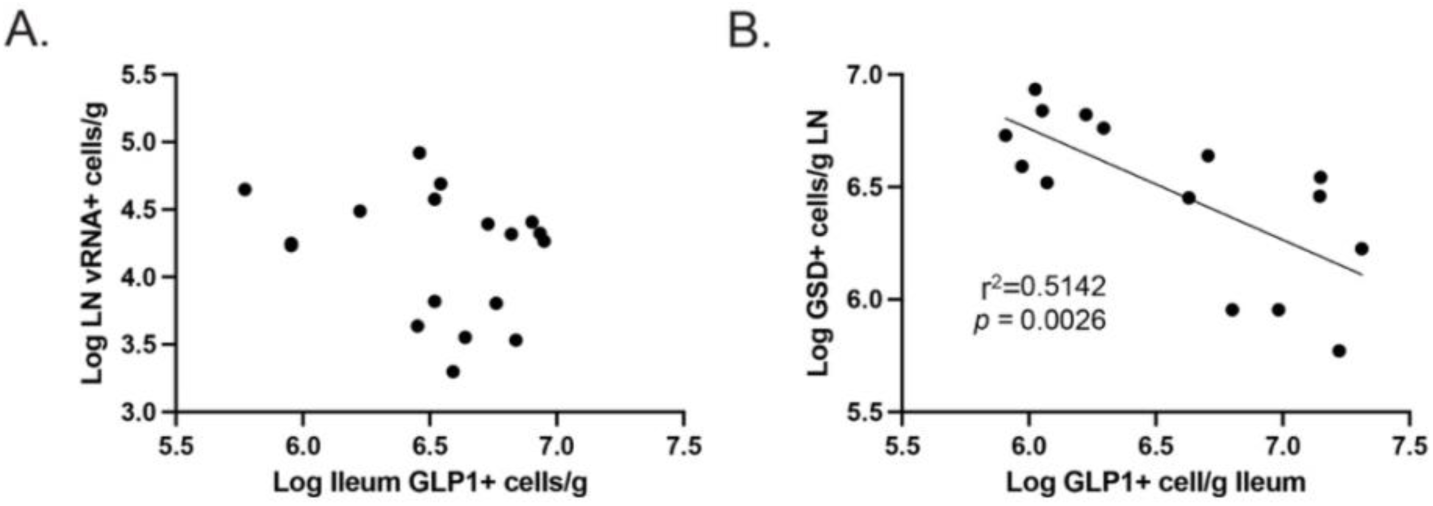
(**A**), Relationship between GLP-1^+^ cells in ileum and vRNA^+^ cells in LNs. (**B**), Relationship between GLP-1^+^ cells in ileum and GSD^+^ cells in LNs.

## Discussion

Our original hypothesis was that ongoing virus production in lymphoid tissues was the direct cause of an increase in proinflammatory cytokines and activated cells, which are common in people with treated HIV infection. Furthermore, we hypothesized that chronic inflammation contributes to the metabolic syndromes that are becoming increasingly common in PWH. Surprisingly, except for plasma measures of IL-8 and cell associated TGF-β, we found no association among proinflammatory cytokines, activated cells, or the cell-associated cytokine production of IL-6 and vRNA^+^ cells, a gold standard measure of HIV production in the LN. While we did find a correlation between the frequency of vRNA^+^ cells in LNs and two biomarkers of immune reconstitution (peripheral CD8^+^ T cells and CD4/CD8 ratio), our findings suggest that an additional mechanism underlies the relationship between HIV production, IA, and downstream cardiometabolic abnormalities in PWH despite effective ART.

Our findings provide the first demonstration of a significant positive correlation between the frequency of HIV RNA^+^ cells and the frequency of cells in the lymph node undergoing pyroptosis. Furthermore, the frequency of GSD^+^ cells correlated with plasma levels of multiple proinflammatory cytokines linked to innate immunity. This observation provides insight into the metabolic syndromes that develop in PWH on long-term ART. Pyroptosis is associated with the development of type 2 diabetes, MASLD, fat depot redistribution, lipodystrophy, and neurodegenerative diseases including Alzheimer disease^24^ ^32^. Also, recent work has suggested that the inflammasome is implicated in HIV-related metabolic syndrome^31^.

Because of the inherent anti-inflammatory properties of GLP-1, we hypothesized that the chronic inflammatory state induced by ongoing pyroptosis would trigger a compensatory increase in the incretin hormone GLP-1^33,34,36,37^. Indeed, we found that (i) GLP-1^+^ cells in the ileum of PWH were increased, relative to HIV-negative controls and (ii) there were significant and direct correlations between GLP-1^+^ ileal cells and many of the proinflammatory cytokines we measured in plasma from PWH. Importantly, however, there was no correlation between HIV vRNA^+^ LN cells and the number of GLP-1^+^ ileal cells. These results suggest that ileal GLP-1 production is promoted by LN virus production but is more directly triggered by innate immune-associated pyroptosis. Because HIV RNA production in LNs correlates directly with the number of pyroptotic cells, which are inversely correlated with GLP-1^+^ ileal cells, these results strongly suggest that HIV production alone is not sufficient to drive ileal GLP-1 production, and that IA-associated pyroptosis drives a counterregulatory GLP-1 response. Taken together, HIV RNA^+^ cell-induced IA, accompanied by APC-derived proinflammatory cytokine induction—a core mechanism in metabolic syndrome pathogenesis—engages the production of the incretin hormone GLP-1 in the gut, which in turn may dampen pyroptosis. Another, non-mutually exclusive potential explanation is the eventual exhaustion of GLP-1-producing cells with increasing pyroptosis. Both possibilities raise the question of whether therapy with GLP-1 agonists, currently in widespread clinical use for metabolic syndrome and type 2 diabetes, may be beneficial by modulating inflammation and pyroptosis in ART-treated PWH with signatures of IA.

This study has limitations. Measures of cytokines and biomarkers were obtained at discrete points in time, and not longitudinally. The mechanisms underlying IA and metabolic comorbidities in PWH likely occur through multiple parallel pathogenic cascades. Therefore, future studies are warranted to measure dynamic responses of circulating GLP-1 levels and to better understand the physiologic relevance of incretins in HIV-related IA.. Nonetheless, this study provides the first data to directly link mediators of chronic IA in HIV pathogenesis to inflammatory cell death and enteroendocrine regulators, providing insight into potential causes of metabolic syndrome, which is becoming increasingly important in PWH on long-term ART.

## Methods

### Sex as a biological variable

Sex was considered a biological variable. Our protocol was written to include any eligible participant regardless of sex, gender, race, or ethnicity, and our recruitment efforts followed that principle. The participant population of this study generally reflected the demographic characteristics of PWH in Minnesota. Most study participants were male.

### Study design and setting

The study was conducted at the University of Minnesota (UMN) and the Hennepin Healthcare Research Institute (HHRI). The main study involved PWH; the first participant was enrolled on October 21, 2021, and the last participant on October 11, 2023. For laboratory comparisons, a cohort of HIV-negative participants was enrolled between October 16, 2023 and July 2, 2025.

### Study participants

Study participants were recruited using institutional review board (IRB)-approved study-specific online and print advertisements posted in local magazines and institutions. Potential participants were also referred from local clinics. Inclusion criteria included age ≥18 years, laboratory-confirmed HIV-1 infection per medical records, stable and consistent ART for over 12 months, plasma HIV RNA <48 copies/mL at screening and over the past 12 months (isolated single blips up to 200 copies/mL were allowed if preceded and followed by undetectable viral load determinations), and screening laboratory tests (complete blood cell count and metabolic panel) within institutional normal range. Exclusion criteria included pregnancy or breastfeeding, and not being a suitable candidate for an inguinal LN biopsy (e.g., current use of anticoagulants, BMI ≥ 30 kg/m^2^, ≥3 LN biopsies in the past). For the HIV-negative cohort, the selection criteria were similar except for the HIV serostatus.

### Study activities and procedures

Screening visits were conducted at either the UMN Phase 1 Clinical Research Unit (CRU) located at the Phillips-Wangensteen building of the UMN Minneapolis campus or HHRI. Other study visits involving data collection, physical exams, blood draws, and procedures took place at UMN only. After successful screening, blood was drawn at a single study timepoint for CBC, metabolic panel, plasma HIV RNA, T cell subset panel, and research experiments at the Schacker Lab and Klatt Lab. Blood was collected in acid citrate dextrose solution A (ACD-A) tubes (Becton Dickinson Inc, Franklin Lakes, NJ, USA) and EDTA tubes. Excisional inguinal LN biopsies were completed by experienced surgeons at the UMN CRU. Ultrasound guidance was used to visualize a potential lymph node, allowing for a more successful and safer intervention. The groin area was scrubbed with an antiseptic solution, and local anesthetics were injected to numb the inguinal area. A 1-inch skin incision was made over the lymph node. After the lymph node was removed (if found), the incision was closed with dissolvable stitches. After completion of the procedure, participants were observed by the study team for a minimum of 2 hours before discharge. Oral acetaminophen was offered post-procedure. Colonoscopies were conducted at the Endoscopy Center of the UMN Medical Center (UMMC) under mild-to-moderate sedation and after overnight bowel preparation. Pinch biopsies were done in the terminal ileum (Radial Jaw 4 Biopsy Forceps, Jumbo with Needle, 3.2 mm; Boston Scientific, Marlborough, MA, USA).

### Clinical laboratory tests

All clinical tests were run in the Clinical Laboratory Improvement Amendments (CLIA)-certified UMMC/Fairview Laboratory. Plasma HIV RNA was measured using the COBAS AmpliPrep/COBAS TaqMan HIV-1 test v2.0 (Roche, Branchburg, NJ, USA. Plasma CD4^+^ and CD8^+^ T cell counts were measured by flow cytometry using the FACSCount™ system (Becton Dickinson Inc, Franklin Lakes, NJ, USA). Both platforms were registered on an external quality assurance program provided by the American Pathologists and Virology Quality Assurance from Rush University Medical Center. The results were shared with the participants and their healthcare providers.

### PBMC and plasma isolation

Tubes were spun at 400*g* for 10 minutes with the brake off. During this spin, 13 ml of Histopaque-1077 (MilliporeSigma, Burlington, MA, USA) was added to each of the SepMate 50 ml tubes (STEMCELL Technologies, Vancouver, Canada). The plasma was removed from the spun ACD-A tubes and frozen in 1 ml aliquots. The plasma was replaced with 1×PBS and transferred to 50 ml tubes, where the volume was adjusted to 32.5 ml. The blood/PBS mixture was pipetted into the prepared SepMate tubes. The tubes were then centrifuged at 1200*g* for 10 minutes at room temperature with a brake. The buffy coat layer was then poured off and divided between two 50 ml conical tubes. Complete RPMI (CRPMI) was added to each tube to bring total volume to 50 ml. Tubes were centrifuged at 400*g* for 10 minutes at room temperature and supernatant was discarded. Pellets were combined and resuspended in 25 ml of CRPMI and then centrifuged at 250*g* for 10 minutes to remove platelets. The pellet was resuspended in 20 ml of CRPMI. A 10 µl aliquot was removed and added to 90 µl of trypan blue for counting by hemocytometer. The cell suspension was centrifuged at 400*g* for 5 minutes at room temperature. The cell pellet was then resuspended in Freezing Media (90% FBS:10% DMSO) to achieve a concentration of 1×10^7^ cells/ml.

### In situ hybridization (ISH)

These methods have been previously described^10,21^. Five-to-ten 5 µm sections separated by 20 µm were analyzed by RNAscope 2.5 (ACD). The anti-sense HIV probes used for the detection of vRNA covers ~4.5kb of the genome and are designed to bind to sequences in *gag*, *pol*, *vif, vpx* (for SIV), *vpr, tat, rev, vpu* (for HIV), *env*, and *nef*. HIV RNA-specific probes from Advanced Cell Diagnostics were used for HIV RNA ISH to identify Clade B viruses (catalog number 416111 and DNA sense catalog number 425531).

### IHC

Five-µm thick sections of paraffin-embedded tissue mounted on glass, positively-charged slides (Creative Waste Solutions) were used for all staining protocols. The tissues were deparaffinized in xylenes followed by graded ethanol and hydrated in deionized water. After antigen retrieval, the tissues were blocked with Sniper (Biocare Medical) for 30 minutes. Primary antibodies were added (**Supplementary Table 3**) and tissues incubated overnight at 4°C. Secondary HRP antibodies (GBI) were added after washing three times in Tris-buffered saline with Tween (TBST). ImmPACT 3,3′-diaminobenzidine (DAB, Vector Laboratories) was added per instructions and counterstained with CAT Hematoxylin (Biocare Medical).

### Quantitative image analysis

Photographic images were captured using brightfield whole-tissue scanning with an Olympus VS200 (Evident Scientific, Inc., Waltham, MA, USA) at 20x/0.80NA. The frequency of vRNA^+^ cells were measured and expressed as the total per unit area, as previously published^38–40^. Briefly, the area of the tissue is measured using image analysis software and expressed as µm^2^. The tissue is 5 µm thick, and by knowing the area of tissue in µm^2^, we then convert that to cm^3^. The average density of lymphoid tissue is 1 gram/cm^3^ providing a measure of cells/g^41^.

High-resolution microscopy was done with the Nikon AX/AX R, Nikon Spatial Array Confocal (NSPARC) system running the Nikon NIS-Elements v6.10.02 with a Plan Apo 60x, 1.42 n.a. oil immersion objective and NSPARC detector with 1.66x zoom was used for sequential collections using the 405 nm laser, 430–475em (DAPI), 488 laser, 525–550em (Atto520), 561nm laser, 570–618em (CF570) and 641nm laser 666–732em (Atto680). Images were tiled using Large Image in ND acquisition. Additionally, the light microscopy data were deconvolved using the Nikon 2D NSPARC automatic deconvolution mode.

### Plasma cytokine and biomarker measurements

Cytokine concentrations were measured in plasma from EDTA tubes using a 13-plex (IL-1β, IL-2, IL-5, IL-6, IL-7, IL-8, IL-10, IL-12p70, IL-17A, IL-23, TNF-α, IFN-γ, and MIP-1β) Luminex panel (Cat. HSTCMAG-28SK, MILLIPLEX) and by ELISA (IL-18) (Cat. DY318-05, R&D systems). All processing was performed according to the kit manufacturer’s instructions. Values below the limit of detection for each analyte were replaced with the lowest detectable concentration divided by two.

### Data collection and statistics

Demographics, medical history, and clinical laboratory results were all collected in a REDCap database. Laboratory results were entered in Microsoft Excel and the Schacker institutional database. Statistical analyses were performed using Prism v10 (GraphPad Software, La Jolla, CA, USA). Participant characteristics and measures were summarized by count (percentage) for categorical variables and by median (IQR) for continuous variables. We used a Mann-Whitney *U* test to compare research lab analyses between groups. *P* values < 0.05 were considered statistically significant.

### Study approval

The UMN IRB approved the study (STUDY00009216). All participants gave written informed consent using IRB-approved forms.

## Data availability

Deidentified data are available from the corresponding author upon reasonable request.

## Author contributions

All authors made substantial contributions to this work. Jo.R. and K.E. provided oversight for the clinical components of the trial at UMN. J.G.C. and G.J.B. performed the LN biopsies. P.A.C., R.C., K.E., J.A., G.W., A.J., Ja.R. C.M.B, and N.R.K. conducted laboratory assays or provided statistical data analysis and interpretation. K.E. and A.A. oversaw administrative details and implementation aspects of the trial and contributed to identifying participants and following them through the clinical protocol. A.K. performed the research colonoscopies. N.K. assisted with some of the blood draws from participants. J.V.B. provided oversight for the conduct of the trial at HHRI. A.T.H. and T.W.S. designed the clinical protocol and provided oversight for the conduct of the trial. P.A.C., K.E., and T.W.S. prepared the manuscript.

## Acknowledgments

High-resolution microscopy was supported by the resources and staff at the UMN University Imaging Centers (UIC), Nikon Center of Excellence. SCR_020997. We would like to thank Amanda Kabage for her help with some of the blood draws, and the staff of the UMN CRU and the study participants who made this study possible. We would further like to thank the staff the UMN Pre-Clinical Research Center for performing Luminex cytokine measurements.

