## Supplementary material for "Ongoing lymphoid HIV production drives pyroptosis and GLP-1 counter-regulation in ART-suppressed infection"

**Supplementary Table 1.** $r^2$ and *p* values for the association between vRNA+ cells in LNs and plasma cytokines that were not statistically significant.

| Cytokine or biomarker (plasma) | HIV RNA+ cells in LNs | |
|---|---|---|
| | $r^2$ | *p* |
| TNF-α | 0.006 | 0.758 |
| MIP-1β | 0.984 | 0.205 |
| IL-23 | 0.019 | 0.581 |
| IL-2 | 0.002 | 0.844 |
| IL-17 | 0.012 | 0.669 |
| IL-12p70 | 0.000 | 0.947 |
| IFN-γ | 0.008 | 0.718 |
| IL-1β | 0.030 | 0.534 |
| IL-10 | 0.129 | 0.156 |
| IL-6 | 0.023 | 0.533 |
| IL-18 | 0.004 | 0.791 |

**Supplementary Table 2.** Relationship between GLP-1+ ileal cells and plasma cytokines. The $r^2$ and *p* values are derived from linear regression.

| Plasma cytokine | GLP-1+ ileal cells | |
|---|---|---|
| | *r* | *p* |
| IL-10* | 0.387 | 0.004 |
| IL-1β* | 0.421 | 0.003 |
| IL-6* | 0.369 | 0.006 |
| IL-18* | 0.293 | 0.02 |
| IL-5 | 0.206 | 0.05 |
| IL-8* | 0.207 | 0.05 |
| IL-23 | 0.182 | 0.07 |
| IL-2 | 0.166 | 0.08 |
| IL-17A* | 0.157 | 0.09 |
| IL-7 | 0.143 | 0.11 |
| IL-12p70 | 0.121 | 0.14 |
| IFN-γ | 0.109 | 0.17 |
| MIP-1β | 0.006 | 0.75 |
| TNF-α* | 0.00 | 0.95 |

*Cytokines associated with pyroptosis.

**Supplementary Table 3.** Antibodies used for IHC.

| **Antibody** | **Company** | **Catalog number** | **Dilution** |
|---|---|---|---|
| GLP-1 | Abcam | ab108443 | 1:100 |
| GSD | Cell Signaling | 36425 | 1:200 |
| IL-6 | Proteintech | 21865-1-AP | 1:1500 |
| CD68 | Biocare Medical | CM033 | 1:400 |
| IL-18 | Abcam | ab191152 | 1:100 |
